# Cooperation of partially-transformed clones: an invisible force behind the early stages of carcinogenesis

**DOI:** 10.1101/431478

**Authors:** Alessandro Esposito

## Abstract

Most tumours exhibit significant heterogeneity and are best described as communities of cellular populations competing for resources. Growing experimental evidence also suggests, however, that cooperation between cancer clones is important as well for the maintenance of tumour heterogeneity and tumour progression. However, a role for cell communication during the earliest steps in oncogenesis is not well characterised despite its vital importance in normal tissue and clinically manifest tumours. By modelling the interaction between the mutational process and cell-to-cell communication in three-dimensional tissue architecture, we show that non-cell-autonomous mechanisms of carcinogenesis could support and accelerate pre-cancerous clonal expansion through the cooperation of different, non- or partially- transformed mutants. We predict the existence of a ‘cell-autonomous time-horizon’, a time before which cooperation between cell-to-cell communication and DNA mutations might be one of the most fundamental forces shaping the early stages of oncogenesis. The understanding of this process could shed new light on the mechanisms leading to clinically manifest cancers.

## Introduction

The cooperation between tumour cells and its environment and the competition between different tumour clones during carcinogenesis are well-established^1^. Other types of cooperations, for instance, the positive cooperation between tumour clones, or even non-transformed clones, have been increasingly recognised as a possible fundamental driving force in cancer as well^2, 3^. The complexity of all possible clonal interactions, particularly during the late stages of cancer, is, therefore, fostering research aimed to model cancer from an ecological perspective^1, 2, 4^. Competition for resources is one of the driving forces for clonal interaction. However, cell-to-cell communication is an equally fundamental mechanism mediating the interaction of cellular populations through shared diffusible or immobile molecules, such as cytokine or metabolites. After early modelling work on angiogenesis^2^, the possibility that partially transformed tumour cells might cooperate was generalised by Axelrod and colleagues^3^. Several recent experimental findings are now supporting the notion that cooperation of clones and polyclonality play an important role in the emergence of cancer.

Glioblastoma multiforme tumours, for instance, exhibit considerable intra-tumoral heterogeneity including the pathogenic expression of an oncogenic truncation of the epidermal growth factor receptor (ΔEGFR) gene and EGFR amplification^5^. The less frequent ΔEGFR clones can support an increased fitness of the more prevalent cells overexpressing EGFR, through secretion of IL6 and LIF and a paracrine effect. Recently, Reeves and colleagues^6^ have used multi-colour lineage tracing with a Confetti mouse line together with the topical administration of a carcinogen, to study clonal evolution during early oncogenesis. Interestingly, the authors observed benign papillomas harbouring an HRAS Q61L mutation with streaks of Notch mutant clones. Although these Notch mutants were considered infiltrating clones with no active role in the oncogenic process, Janiszewska and Polyak^7^ noted that cooperation between the Notch and HRAS mutants could not be excluded and that streaks of Notch clones are reminiscent of structures found in non-mutualistic colonies of budding yeast. Although unproven, it is conceivable that the less frequent clones can provide, altruistically, a fitness advantage to the HRAS mutant cells similarly to what has been observed for glioblastoma multiforme^5, 8^ or for WNT-secreting wild-type HRAS clones supporting HRAS mutants^9^. While facilitating the oncogenic process, a non-mutualistic clone would be then outcompeted by more aggressive clones after a clonal sweep and diversification into multiple intermixed mutants^6^ suggestive of mutualistic clonal interactions^7^.

However, it is unclear if these observations, often obtained using model systems with carcinogens or established tumour clones, can be recapitulated at the low mutational rates occurring naturally^10^. Furthermore, it is unknown at which stage of carcinogenesis, non-cell-autonomous mechanisms might have a role^9^. As cell-to-cell communication and clonal interaction are often neglected in formal models of carcinogenesis, we propose a model for the interaction between the mutagenic process and cell-to-cell communication within a three-dimensional tissue architecture. We developed the simplest possible models to capture the basic emergent properties of early oncogenesis in the presence of mutations and clonal cell-to-cell communication. We propose that the extremely low mutational frequency encountered in physiological conditions does not render cooperation between mutations in adjacent cells unlikely but – rather the opposite – that synergy between the mutational process and cell-to-cell communication might play a fundamental role in carcinogenesis.

## Results

### A model for mutationally-driven cooperation in oncogenesis

The question addressed in this work is not *if* cooperation between mutant (partially-transformed) cells can occur, but *how likely* or *when* distinct mutations can occur in different cells cohabiting within the same tissue. Therefore, we develop a simple mathematical model to gain insights into answering these fundamental questions. We consider a low mutational rate ρ_0_, constant throughout oncogenesis and equal for each possible oncogenic mutation^11^. With oncogenic mutation, we refer to any mutation that at any given time (not necessarily when it occurs) might contribute to the increased fitness of a clone that will eventually evolve into cancer, either through cell-autonomous or non-cell-autonomous mechanisms.

The probability for a single cell to accrue two specific mutations independently within a given time interval is thus p_0_^2^ (with p_0_ = ρ _0_ t << 1). The probability that two neighbouring cells exhibit one given mutation each independently is, unsurprisingly, the same. Initially, we assume non-dividing cells in a well-organised tissue that after accumulating these two mutations acquire a fitness advantage. We will refer to these cells as initiated or transformed, but we will use these terms very loosely only to indicate a gain in fitness.

In tissue with N cells, the probability of cell-autonomous initiation of one mutant cell is simply p_ia_=Np_0_^2^p_a_. Similarly, the probability of non-cell-autonomous initiation is p_in_=NCp_0_^2^p_n_. p_a_ and p_n_ are defined as the probabilities that one cell harbouring the right pair of mutations – either by itself or within its neighbourhood – survives tumour-suppressive mechanisms (**Fig. 1a**). C is a coordination number, *i.e.* the number of cells within the neighbourhood of a reference cell (**Fig. 1b, c**). Within the validity of common assumptions (*e.g.*, equally probable, spatiotemporally-invariant and independent mutational events), the probability of initiation within a group of N cell is the sum of p_ia_ and 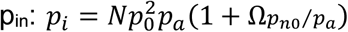 with Ω = *Cp*_*n*_/*p*_*n*0_ and where p_n0_ is the probability that one cell is transformed when directly in contact with another mutant.

**Figure 1.**
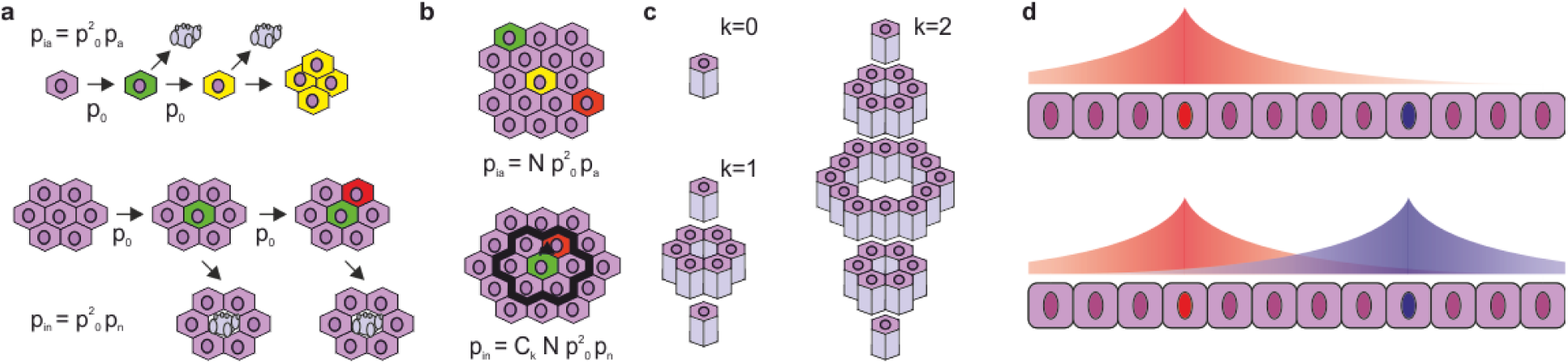
Tissue organisation and non-cell-autonomous mechanisms. **a**) A simple model where cells accumulate two mutations (top) or two mutations occur in different cells within the same neighbourhood (bottom). When the probability to accrue mutations is low, within a tissue of N cells, there will be more opportunities for mutations to co-occur within a given neighbourhood (bottom) rather than within the same cell (top). **c**) The neighbourhood of a cell can be described as a problem of geometrical tessellation of space which will depend on tissue organisation, here shown a simple example of hexagonal pillars tesselating space. **d**) gradients of shared resources (e.g., growth factors or metabolites) might be then induced by either one or the other cell triggering interactions by juxtacrine or paracrine effects.

p_n_ (and thus Ω) depends on tissue organization and the type of cell-to-cell cue that contributes to the process of transformation (**Fig. 1d**). With this simple notation, the answer to our central question can be thus separated into the study of tissue organization (the factor Ω) and the magnitude of p_n0_ compared to p_a_.

### Tissue organisation

To model the organisation of tissue wherein mutated cells are resident, several aspects of tissue organisation have to be considered:

i. the more distant a neighbouring cell is, the lower the probability of cooperative non-cell-autonomous effects should be, *i.e.* p_n_ shall be a function of distance (*d*);
ii. *C* is the sum of cells in extended neighbourhoods or the sum of *C*_*k*_, *i.e.*, the number of cells in the k-neighbourhood (at a distance *d*_*k*_), where k=1 defines cells in contact (*i.e.*, d_1_=0);
iii. *C*_*k*_ depends on tissue architecture that we model as a problem of three-dimensional tessellations of space;
iv. tissues are compartmentalised and, therefore, boundaries effects should be considered.

Therefore, in general, the factor Ω can be described as the cumulative effect on the probability of initiation of a reference from each cell within a tissue as consequence of a cell-to-cell communication, 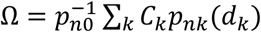. For convenience, we describe *C*_*k*_ just for two different tissue topologies, a tissue organised in stacked hexagonal pillars or a thin layer of similar hexagonal pillars. In the former case, cells tessellate a three-dimensional space, and we neglect effects at the periphery. In other words, we assume that the number of cells contained within a tissue is larger than the cells at its periphery. **Fig. 1** illustrates the progression of the number of cells included in subsequent neighbourhoods. In **Appendix 1** we demonstrate that C_k_ = 6k^2^+2. For a significantly more constrained topology where only three layers exist *C*_1_ = *s*_0_ + 2 and *C*_*k*>1_ = *s*_0_(3*k* − 2). This description permits us to evaluate analytically the effects of tissue organisation on the probability of cooperation between mutations. In the next section, we provide also numerical examples showing the general validity of this model in the presence of even more stringent topological constrains.

### Oncogenic field effect

Without loss of generality, we assume that the interaction between two mutant cells is mediated by a sharing diffusible product^3^, for instance a growth factor or a metabolite. Eldar *et al.* have modelled how the concentration of a signalling molecule (a morphogen) secreted by a cell decays in space^12^. Typically, the morphogen concentration is abated by passive diffusion and linear degradation resulting in exponentially decaying concentration gradients. However, ligand-morphogen interactions can induce non-linear mechanisms of morphogen degradation resulting in power-law decays. Therefore, we first analyse the decay of an oncogenic field akin morphogen gradients using power or exponential decays because of their physiological relevance^12-15^.

For the case of a power function (p_n_(k) = p_n0_ k^-l^) and a three-dimensional tissue described by hexagonal pillars (C_k_ = 6k^2^+2), the factor Ω can be described analytically with Ω(*l*) = 6*ζ*(*l* − 2) + 2*ζ*(*l*) (**Appendix 2**). *ζ* is the Riemmann Zeta function and is finite only for an argument larger than one (here l>3). Therefore, for a large interconnected tissue, oncogenic biochemical gradients induced by a mutant cell must decay very steeply for non-cell-autonomous mechanisms not to dominate. In the limiting case where only the 1-neighbourhood is relevant for transformation (*l* → ∞), the Riemann Zeta function converges to unity and therefore Ω = 8. This is just the number of cells in direct contact to the reference cell (C_1_) showing mathematical consistency and providing a lower boundary to Ω in the case of small effects in a very constrained topology. Conversely, for shallower gradients where the Riemann Zeta function does not converge (l<4), these probabilities will be significantly larger. We obtained these results modelling tissues of non-finite extensions to derive analytical solutions. However, through numerical estimations, it is simple to demonstrate how these observations are generally valid and correct also for small volumes of cells (**Fig. 2a-b**). For example, in a small neighbourhood with a radius of 10 cells, Ω ∼ 11.5 (l=4) and Ω ∼ 340 (l=1), values (**Fig. 2a**, solid circles) that reach 12 and 3 10^4^, respectively, for a neighbourhood with a radius of 100 cells (**Fig. 2a**, empty circles). Similarly, we demonstrate that for a thin three-layer tissue, Ω = 12*ζ*(*l* − 1) − 8*ζ*(*l*) + 4/3 (see **Appendix 2**). This series converge for l>2, it assume a value of 11.7 for l=3, and numerical estimations show that Ω reaches values of ∼24 and ∼50 for l=2 within a limiting neighbourhood with a 10 or 100 cell radius, respectively (**Fig. 2a**, empty and solid circles). In the limit case where only the first neighbourhood is relevant (*l* → ∞), Ω ∼ 5.3. Therefore, even within this rather constrained topology, Ω obtains rather large values.

**Figure 2.**
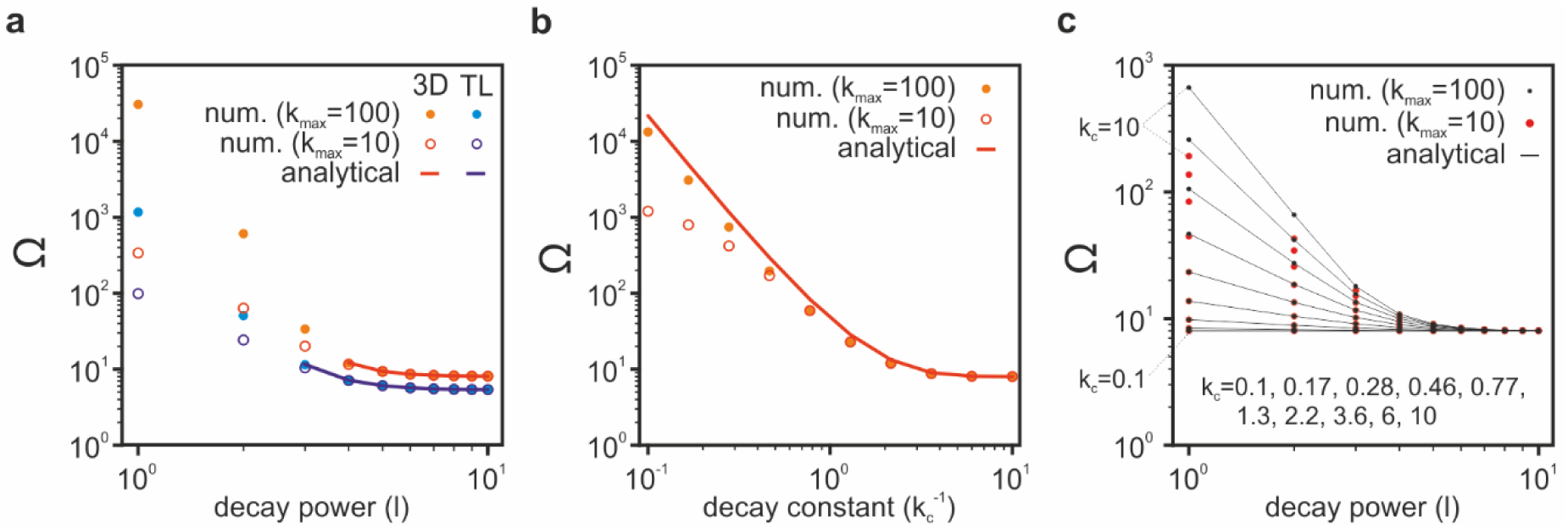
Numerical validation of the oncogenic field solutions. **a**) Comparison between numerical and analytical solutions to estimate the value of the oncogenic field factor Ω for a three-dimensional (3D) and three-layered (TL) tissue model. The solid lines represent the analytical solutions within the limits of its convergence (l>2 for TL in blue and l>3 for 3D in red). The numerical estimations, computed over a small 10 (empty circles) and larger 100 (solid circles) cells radius neighbourhood, confirms the observations we reported, more generally, on the analytical solutions, including the necessity of steep power decays and a convergence to Ω∼8 (3D) and Ω∼5.3 (TL) for these limiting cases. **b**) Identical computations described in a) but for the exponential decay model for a three-dimensional tissue. Both the analytical solution and numerical estimates converge to the value of Ω∼8 for steep decays. **c**) Values for Ω computed for a general case where the oncogenic field decays jointly as the inverse of a power-law and exponentially. The analytical and numerical solutions are shown for the same parameter sweep shown in a) and b), i.e. with the inverse power from 1 to 10 and with a decay constant kc from 0.1 to 10.

Gradients described by power functions are shallower than exponentially decaying gradients at longer distances. Although both gradients are physiologically relevant, power-like functions might overestimate Ω. The formalism for exponentially decaying oncogenic fields is less elegant (see **Appendix 3**); however, it can be readily demonstrated that even for steep gradients decaying of a third at every cell distance (k_c_=1), Ω can assume double-digit values (see also **Fig. 2b**).

The analysis of power-law and exponential decays are rather instructive, and they are often used to model morphogen gradients as a solution to the reaction-diffusion equation for one-dimensional problems and for specific three-dimensional architectures. We can also demonstrate (**Appendix 4**) that for 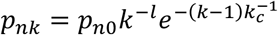, *i.e.*, when the oncogenic gradients jointly decays as an inverse power-law and exponentially, 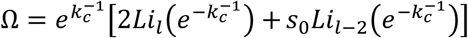, where *Li* is the polylogarithm. This analytical solution describes the expectation for Ω for an oncogenic field induced by stochastic (mutationally-driven) point-sources of shared resources in an ideal three-dimensional tissue (see **Fig. 2c**) in the presence of degradation. Once again, at the limit for a fast decaying concentration gradient, the value of Ω∼8, long-distance interactions (k_c_>>0) can drastically increase the magnitude of Ω and with high values found also for small clusters of cells (**Fig. 2c,** blue and red circles).

We can thus infer a general consideration from the mathematical description of the proposed case studies that are aimed to exemplify the possible synergy between the mutational process and non-cell-autonomous effects. Unsurprisingly, the specific tissue geometries and the properties of concentration gradients results in rather different magnitudes of an ‘oncogenic field’. However, either through *juxtacrine* (contact-dependent) or *paracrine* (short-or long-distance) signalling, mutations in tissue neighbourhoods that can cooperate through cell-to-cell communication are likely to have a significant role in oncogenesis, in addition to mutations co-occurring within a cell.

### Cell-autonomous time-horizon

So far, we have discussed *if* and *how likely* mutationally-driven non-cell-autonomous mechanisms might be; next, we address the question about *when* these mechanisms are more likely to occur. Indeed, we have shown that non-cell-autonomous mechanisms can increase the probability that mutations contribute to carcinogenesis by a factor Ω. A corollary to this observation is that cooperation between non-transformed cells might contribute to tumour initiation earlier than cell-autonomous mechanisms. For simplicity, we consider only the mutational process and neglect p_a_, p_n_ and p_n0_. One cell accrues pairs of mutations at the rate *ρ*_0_ but within a neighbourhood cooperating cells accrue mutations at an apparent rate of 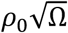. We tested this simple mathematical inference with Monte Carlo simulations (see **Fig. 3** and **Methods**). We simulate the independent and stochastic appearance of four types of mutations (A, B, C and D) at a rate of *ρ*_0_=10^−6^ mutations/day on a lattice of N=10^6^ cells with 2,000 replicates. When one cell accrues mutations A and B, it is flagged as an AB mutant; when a cell becomes a C mutant and in its neighbourhood there is a D mutant, the C mutant will be listed as a CD (cooperating) clone. The average time for a double-mutant cell to appear (<t_AB_>) is ∼29 months (**Fig. 3a**). At the net of noise, the distribution of <t_AB_> values (**Fig. 3a**, grey curves) depends only on *ρ*_0_ and N but not on Ω and, therefore, we overlay the average of the four distributions of <t_AB_> values (**Fig. 3a**, black curve) as reference for the cooperating mutants. The distribution of the waiting times for the appearance of CD clones depends on the value of Ω (**Fig. 3a**, coloured curve) and exhibits an average time (<t_CD_>) of about 14.7, 10.1, 8.5 and 6.2 months for Ω values equal to 4, 8, 12 and 24, respectively – scaling as Ω^−0.5^. Thus, we can define *t*_*a*_ (<t_AB_> in our simulations) as the average time for a tissue of N cells to accrue two mutations which is inversely proportional to *ρ*_0_N. By definition, *t*_*a*_ -the time-horizon after which cell-autonomous mechanisms might dominate -is preceded by a latency period during which single mutations are more likely. However, our model predicts the existence of a significantly long period *t*_Ω_ = *t*_*a*_Ω^−0.5^ ≤ *t* < *t*_*a*_ when mutationally-driven cooperation between adjacent cells is more likely than mutationally-driven cell-autonomous mechanisms to occur. For instance, in the limiting case where only the first-neighbourhood significantly contribute to tumour initiation (Ω = 8), during ∼65% of the time interval preceding *t*_*a*_, clonal cooperation is likely to be a fundamental mechanism that synergizes with the mutational process to support partially transformed clones. For Ω values of 4, 12 and 24, this interval will be about 50%, 70% and 80% of *t*_*a*_. The scaling of *t*Ω as *t*_*a*_Ω^−0.5^ is shown in **Fig. 3b** from the plot of <t_AB_>/<t_CD_> (**Fig. 3b**, red) and the scaling of the number of mutation in a given neighbourhood defined by Ω at *t*_*a*_ is shown is <N_CD_> (**Fig. 3b**, blue). Because of the stochastic nature of the mutational process, the distribution of waiting times for double mutants (AB and CD) are broad. This heterogeneity result in no CD co-operating clones in about 40%, 20%, 15% and 8% of the simulation runs with Ω = 4, 8, 12 and 24, respectively. The majority of the cases, however, co-operating mutants preceded AB-clones and for larger values of Ω the probability for mutations to appear within a tissue neighbourhood is so high that CD mutants are likely to reoccur multiple times randomly (**Fig. 3c**). Therefore, even assuming a role for synergy between the mutational process and cell-to-cell communications during the earliest steps in oncogenesis (*t* < *t*_*a*_), chance will determine the first occurrence of co-or cooperating-mutations (as a function of Ω), possibly influencing the evolutionary trajectory of a tumour and contributing to tumour heterogeneity.

**Figure 3.**
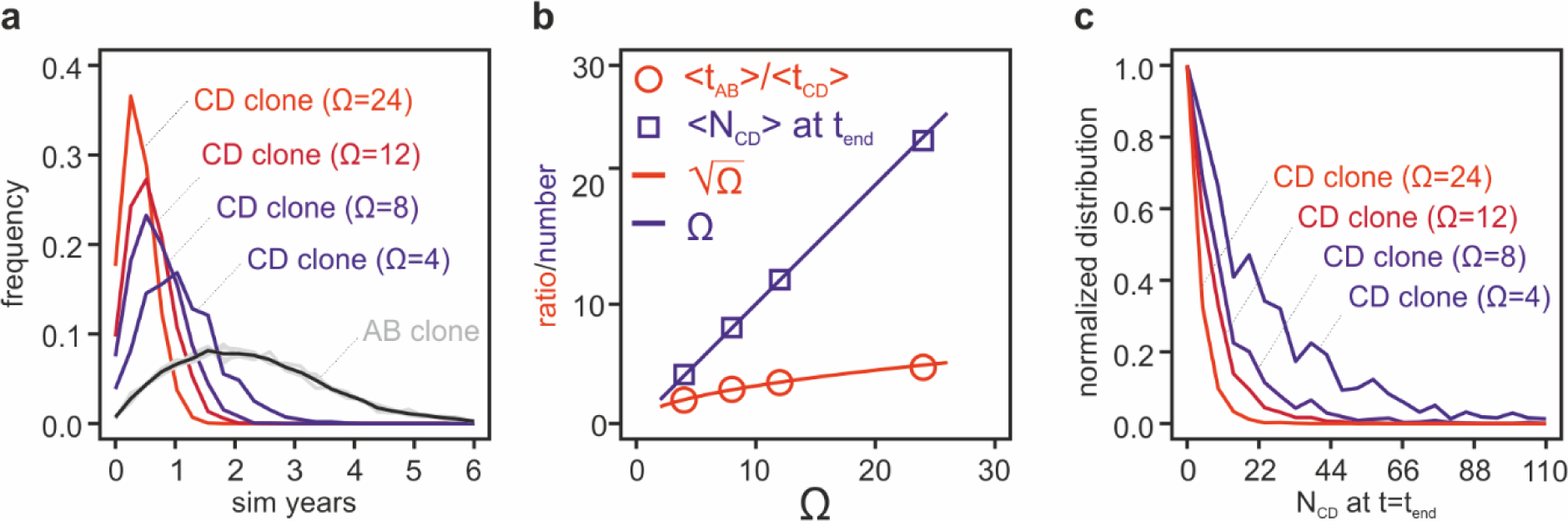
Monte Carlo simulations of the cell-autonomous time-horizon. **a**) Probability distribution of the waiting-times for the occurrence of the first two co-occurring mutations (AB-clones, grey and black curves) or for the first cooperating mutations (CD-clones, coloured curves) through non-cell-autonomous mechanisms for Ω values equal to 4, 8, 12 and 24. The grey curves are the four independent simulation runs related to these four conditions and are identical except for noise; the black curve is the average of these four runs and can be used as reference. **b**) The average time (<t_CD_>) at which the first cooperating CD-clone is observed scales with the square root of Ω (red line) compared to the average time (<t_AB_>) at which the first AB-mutant appears. The average number of cooperating mutations within a neighbourhood at t=t_end_ (<N_CD_>) scales as Ω (blue curve). **c**) Distribution (normalized to maximum for better visualization) of the number of CD-clones at the end of the simulations (t=t_end_). t_end_ is the time at which at least one AB-and one CD-clone are detected.

## Discussion

A role for non-cell-autonomous mechanisms in cancer is well-established, often as a mechanism of interaction between cancer cells and the surrounding tissue^2, 16-19^. The cooperation of non-or partially-transformed clones as a driving force underlying oncogenesis has also been hypothesized^3^, and there is nowadays accumulating evidence suggesting that a description of oncogenesis focused exclusively on cell-autonomous mechanisms might under-represent the importance of oncogenic signalling in cancer^5, 9, 19^.

Experiments in *Drosophila melanogaster* have also shown that inter-clonal cooperation between mutants harbouring an oncogenic KRAS mutation or inactivation of the tumour suppressor *scrib* can support tumorigenesis mediated by JNK and JAK/STAT signalling^20^. Recently, Marusyk and colleagues (2014) have used a mouse xenograft model to test the effects of clonal heterogeneity demonstrating that clones expressing the chemokines IL11 are capable of stimulating overall tumour growth through a non-cell-autonomous mechanism, while clonal interference maintains genetic intra-tumour heterogeneity^9^. Similarly, Inda and colleagues (2010) have shown how intra-tumour heterogeneity observed in glioblastoma can be maintained through cross-talk between mutants harbouring a ΔEGFR that secrete IL6 and LIF to support fitness in clones with EGFR amplification^5^. Clearly and colleagues (2014) has also shown that WNT-producing HRAS wild-type clones can support tumorigenicity and clonal heterogeneity by cooperating with clones harbouring mutant oncogenic HRAS^21^. These observations support the emerging notion that intra-tumoral heterogeneity is often of polyclonal origin and is an active process supported by non-cell-autonomous mechanisms. However, the role for poly-clonality and clonal cooperation during earlier stages of oncogenesis can be seen in contradiction with the low estimates of mutational rates in cancer^10^. Furthermore, it is unclear if clonal cooperation has a role during early oncogenesis or only at later stages when a heterogeneous tumour is established^21^.

Aiming to contribute to filling this gap in knowledge, we developed simple analytical and computational models for mutation-driven oncogenesis in the presence of cell-to-cell communication. Using similar assumptions used to model mutationally-driven oncogenesis, we have studied *if, how* and *when* is likely that cell-to-cell communication might cooperate with the mutational process from the perspective of basic principles. Our analysis raises provoking observations on the earliest steps in oncogenesis. We show that irrespectively of the background mutational rate if a set of transforming mutations are sufficiently likely to occur within a single cell in the lifetime of a patient, an equally rare yet oncogenic set of mutations are equally (or more) likely to contribute to tumorigenesis through non-cell-autonomous mechanisms. We have introduced the parameter Ω which capture the impact of tissue organisation and non-cell-autonomous mechanisms on cancer evolution. We modelled non-cell-autonomous mechanisms in analogy to morphogens during embryonic developments. Ω describes the magnitude with which paracrine, juxtacrine and other mechanisms mediated by shared substrates (*e.g.*, growth factors and metabolites) might impact the transformation of a cell or clone. As such, Ω represents an oncogenic field effect, where oncogenic fields have the opposite outcome of morphogens by contributing to the de-regulation of tissue homeostasis. Furthermore, we have identified a stage of oncogenesis during which clonal cooperation might not simply coexist with clonal competition but even dominate before the emergence of clones capable of growing autonomously. With the help of our model, experimentally, the problem is reduced to the measurement of quantities such as p_n0_ and p_a_ or the abundance of genes that, once mutated, can drive oncogenesis by non-cell-autonomous mechanisms. We argue that the magnitude of the oncogenic field effect (Ω) and the prediction of an autonomous time-horizon suggest a significant role for mutationally-driven and non-cell-autonomous mediated poly-clonal evolution of cancer during, at least, a very early stage of oncogenesis.

The model described here is purposely simple aiming to illustrate the basic principles emerging by the cooperation of the mutational process with non-cell-autonomous mechanisms^3^, a phenomenon that, to our knowledge, is often neglected when models of somatic evolution of cancer are studied analytically^22^. For this reason, we did not include the description of more complex and important features of real tissues such as clonal dynamics, tissue homeostasis, tissue mechanics, and other mechanisms for gradient formation of biomolecules. Each of these processes can change considerably the magnitude of the effects we described. The concentration gradients on their own, for instance, can be enhanced by compartmentalisation, abrogated by diffusion into lumens or the vascular system, or affected by systemic alterations of shared resources (*e.g.*, hormones, lipids). If a proliferative tissue is considered, with a fitness advantage for cooperative clones compared to wild-type cells, the presence of these non-or partially-transformed clones could be even more significant, increasing the probability to accrue further mutations at a faster pace and shaping the initial period of oncogenesis.

However, tissues are complex systems and diverse mechanisms of tissue homeostasis in different tissues might conflict with this perspective. In the case of a fast self-renewing tissue like the intestinal epithelium within which the cell-of-origin for common tumours is likely to be a stem cell^23^, the highly compartmentalized stem-cell niche might pose an effective barrier to oncogenic field effects. The intestinal epithelium is one of the most proliferative tissues subject to a high mutagenic burden and it has been broadly studied both mathematically^24, 25^ and experimentally^23, 26, 27^. A small group of adult stem cells reside within the colonic crypt and maintain the homeostasis of the villi lining the intestine^23^. Within the crypt and the villus, the balance between proliferation and differentiation is maintained by a complex network of signals (*e.g.*, WNT, NOTCH, BMP, and EGF) generated by specialized Paneth cells within the crypt and cells within mesenchyme lining the crypt^27^. Mutations in the WNT (*e.g.*, APC or CTNNB), EGFR (*e.g.*, KRAS, PIK3C or BRAF) and TGF-β (*e.g.*, SMAD4) signalling pathways gradually render cells independent from niche signals to grow autonomously and promoting cancer. While it is still more likely that two mutations are acquired within adjacent stem cells rather than within one cell, neutral genetic drift or selection fix or purge genetic mutations within the crypt that will be thus, most of the times, monoclonal^23^. However, not all mutations occurs in the stem cells within the crypt^28^ and tissue homeostasis is maintained by crypt fusion and fission leading to field cancerization and the possibility that partially-or non-transformed clones might interacts not within a monoclonal crypt but between patches harbouring different mutations^23, 29^. These considerations might hold true also for other highly proliferative tissues, such as the well-described human epidermis and the oesophageal epithelium^30-33^. In these tissues homeostasis is maintained by a balance between the probability for progenitor cells to divide symmetrically or asymmetrically, giving birth to two progenitor cells, two differentiating cells, or – more commonly – one differentiating and one progenitor cell^30, 33^. Alcolea and colleagues, for example, have shown that mutations in the NOTCH pathway reduce the probability for a progenitor cell to generate two differentiating cells and induce wild-type cells to differentiate^30^. In combination with P53 mutations, this cell fate imbalance leads to field cancerization^30^. Considering the experimental observations on later stages of carcinogenesis we have already discussed^5, 6, 8, 9, 20, 21^ and our results, it is conceivable that mutations might also cause cell fate imbalance through non-cell-autonomous mechanisms during early oncogenesis. As cell fate determination and the occurrence of mutations are stochastic processes, the role of non-cell-autonomous mechanisms not only might vary across different tissues depending on their organization but also within the same tissue of origin. We described in Fig. 3 that the occurrence of a double-mutant clone that might acquire a fitness advantage autonomously, even if less likely, can still precede the occurrence of two cooperating single-mutants. Similarly, genetic drift and selective pressure could either maintain or collapse one or both of the cooperating populations^9^. Therefore, each tissue and each tumour might be affected differently by non-cell-autonomous mechanisms, mechanisms that could alter the evolutionary trajectory of tumours that later acquire full independence from cooperating clones thus contributing also to tumour heterogeneity. As computational modelling of multicellular tissues can describe complex homotypic and heterotypic interactions, including short-and long-range interactions, and tissue mechanics^34-36^, computational models rather than analytical tools might be more appropriate to investigate the possible role for ‘oncogenic fields’ in complex mutagenic environments.

While the mathematical analysis we presented was not elaborated to capture more complex phenomena occurring during oncogenesis, our model highlights the importance of identifying the genes and the shared resources that can mediate clonal cooperation, such as growth factors (*e.g.*, mitogens, interleukins, *etc.*) or even metabolic by-products that are often at the basis of cooperative behaviour in lower organisms^37-39^. The theory described here was aimed to be simple and, at the best of our knowledge, it is a first attempt to describe the problem with an explicit mathematical model to extend the existing models of oncogenesis^11, 22, 40-43^ to non-autonomous mechanisms. Indeed, we emphasise that our model is not in contradiction with the prevailing models of oncogenesis, as it is based on similar assumptions, but it highlights an equally important role for tissue organisation and cell-to-cell communication that cooperate synergistically with a mutationally driven process.

## Methods

The mathematical demonstrations of the analytical equations presented in this work are described in **Appendices 1-3**.

The numerical evaluation of these analytical results (**Fig. 2a-c**) was performed with the Matlab script ‘*analytical_and_numerical_comparisons_v4’* (Mathworks, version 2018) freely available from the GitHub repository *alesposito/CloE-PE*. The numerical estimations simply compare the value obtained from the approximated analytical solutions described in the appendices to direct numerical estimate computed on given neighbourhoods with features described in the main text. For the case of a three-dimensional tissue (**Appendix 2**) the values of the analytical solution Ω = *s*_0_*ζ*(*l* − 2) + 2*ζ*(*l*) were compared to those of the finite series 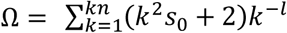 for different power functions. Data shown in **Fig. 2a** are computed with *kn* equal to 50 and 100, *l* ranging from 1 to 10, and *s*_*0*_ = 6. Within the same parameter space, we compare the analytical description of a three-layer thin tissue Ω = 12*ζ*(*l* − 1) − 8*ζ*(*l*) + 4/3 to the numerical estimates of 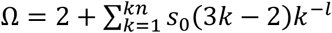. Similarly, **Fig. 2b** shows a comparison between the analytical description 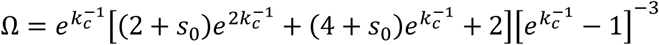 of a three-dimensional tissue with an oncogenic field decaying as an exponential function (**Appendix 3**) and the finite series 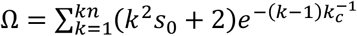. Also in this case, the parameters used in the numerical evaluations were *kn* = 50 and 100, and *s*_*0*_ = 6 with the inverse of the decay constant 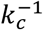 spanning the 0.1 to 10 range. Last, for the distribution jointly decaying as a power and exponential function (**Appendix 4** and **Fig. 2c**), numerical estimates of 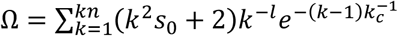 were compared with the analytical solution 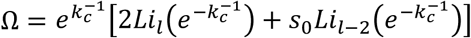 on the same parameter space described for the other cases.

Thee Monte Carlo simulations (**Fig. 3**) used to evaluate the relationship between the time-horizon for cell-autonomous mechanisms (*t*_*a*_) and non-cell-autonomous mechanisms are available as the Matlab script ‘*polyclonal_mutation_cooccurence_check_v4’* freely available from the GitHub repository *alesposito/CloE-PE*. We simulated a lattice of 10^6^ cells with a mutational rate equal to 10^−6^ mutations per cell per simulated day (simday). At each simday and at each node of the lattice, four random numbers (n_m_, with m=A, B, C or D) were drawn from uniformly distributed numbers in the [0,1] interval. For any of the indexes A, B, C, or D where n_m_ was lower than or equal to 10^−6^, the correspondent cell was switched from non-mutant to mutant. Cells were then allowed to accumulate these four mutations for a maximum of 100,000 simdays. When a cell acquires both A and B mutations, an AB-mutant cell is established and logged. When a D mutation appears in a neighbourhood of a C-mutant, a CD-cooperative clone is logged. CD-cooperative mutants are detected utilizing convolution filters that detects the co-occurrence of a D mutant within the centre of a reference neighbourhood and C-mutant in its immediate vicinity. The publicly available code implements the following neighbourhoods scans equivalent to Ω value of 4, 8 and 12. For Ω = 4, detection in position North (N), East (E), South (S) and West (W); for Ω = 8, as for the previous case but with the addition of NE, NW, SE and SW; for Ω = 12, as for the previous case but with the addition of one non-adjacent cell in N, E, S and W position. As soon as at least one AB-and one CD-cooperating clone occurs, the simulation is interrupted. Simulations are then repeated 2,000 times and the distributions of the appearance of first AB-or CD-clones, and number of CD-clones at the appearance of an AB-clone are generated.

The scripts were run on a Dell Precision 5810 workstation utilizing an Intel Xeon E5-1625 CPU and 64GB RAM. The four Monte Carlo simulations shown in Fig. 3 are computationally intensive and were ran in parallel for about two days.

## Acknowledgements

A.E. acknowledges the financial support provided from Medical Research Council program grants (MC_UU_12022/1 and MC_UU_12022/8) awarded to Prof. Ashok Venkitaraman.

## Appendix 1 Description of tissue organization

The 1-neighbourhood of an individual cell will contain 6 adjacent cells within the plane and 2 polar cells, one at the top and one at the bottom of the reference cell. The 2-neighbourhood will contain 12 cells within the plane, 6 cells in the above and bottom layers and two polar cells. The 3-neighbourhood will have 18 cells within the plane, 24, 12, 6 and 2 in the other layers. By induction, we infer that the k-neighbourhood of an individual cell in such tessellation is made of ks_0_ cells within the plane, the sum of is_0_ for each of the above and below layers, with i from 1 to k-1, and the two polar cells:

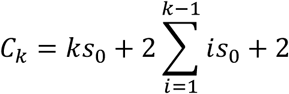

To simplify, *ks*_*0*_ (*s*_*0*_ here is 6) can be brought within the sum, then allowing to simplify to the correspondent triangular number:

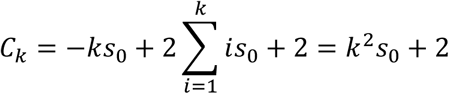

It should be noted that for intercalated layers, the coordination values can be larger.

Another example, this time including a significant constraint in topology, is represented by the same topology, where only three layers are considered.

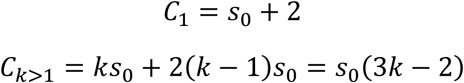

## Appendix 2 Probability of initiation (power function)

Let’s assume the probability of tumour initiation is proportional to a concentration gradient, similar to a morphogen, or an oncogenic mitogen/morphogen field decaying as a power function:

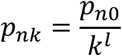

Where *p*_*n0*_ indicate the probability of tumour initiation when cells are attached (the 1-neighbourhood). Therefore, in the case of 3D hexagonal tessellation, we can derive the factor Cp_n_:

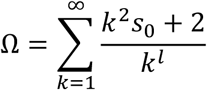

This sum is carried over an infinite neighbourhood, and the validity of the results will be checked numerically. First, we can expand p_tn_:

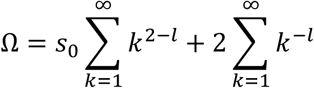

These series can now be described by Riemann Zeta functions:

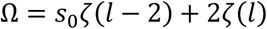

Let’s now consider the thin 3-layer tissue which tessellation was already discussed. In this case, for one cell:

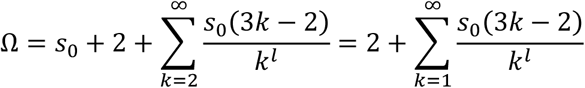

Following the same process described before, we can obtain:

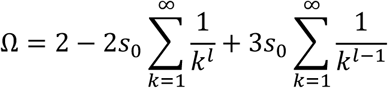

And,

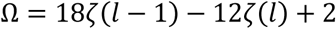

This describe the probability of transformation for a cell in the middle layer. We can approximate the result over the tissue equal to this value by N/3 (middle layer) and with half contribution for the top and bottom layer resulting in

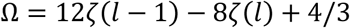

## Appendix 3 Probability of initiation (exponential function)

Let’s now assume the oncogenic field decays as an exponential function:

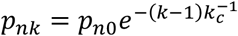

Where *k*_*c*_ is a decay constant expressed in terms of k-neighbourhood for simplicity. If two cells are in contact, the probability of initiation will be p_n0_ as per definition of p_n0_. When cells are at a k_c_+1 distance, this probability is 1/e lower, i.e. ∼30% lower. In the case of 3D hexagonal tessellation, the factor *Cp*_*n*_ can be now expressed as:

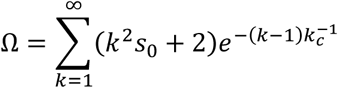

Or the sum of the series:

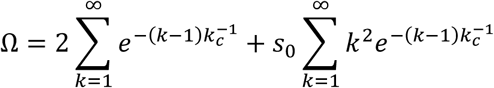

The first series converges to:

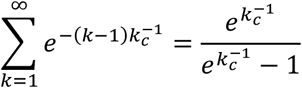

The second series can be represented as:

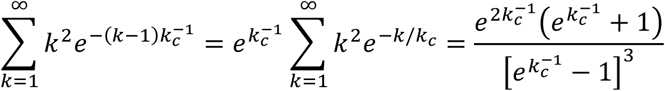

Therefore,

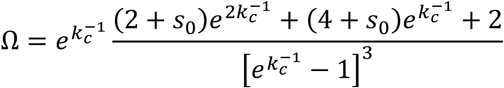

With s_0_ = 6, once again to confirm mathematical consistency, 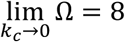, as in the case where only adjacent cells are important. Shallower decays will again increase this value (see Fig. 2).

## Appendix 4 Probability of initiation (generalization)

We have characterized the oncogenic field in relation to typical descriptions of morphogenic gradients^15^. While relevant for specific cases, steady-state concentration gradients of shared resources in space, generated by passive diffusion and linear or non-linear degradation, can adopt different shapes. One useful analytical description is represented by concentrations that decay as the product of exponential and power-law functions, for instance as:

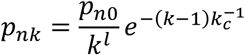

With the same formalism and strategies described in **Appendices 2** and **3** we can show that, for a three-dimensional tissue:

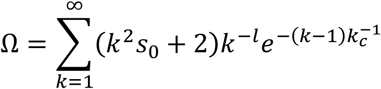

This analytical representation of Ω can be expressed as sums of polylogarithm functions:

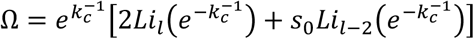

This representation converges to those shown in **Appendix 2** and **3** in the cases where *k*_*c*_ is very large or where *l* is very small, respectively, *i.e.* when the power-law or the exponential decay components are negligible. The case l=1 represents an oncogenic field induced by continuous point-sources in an unconstrained three-dimensional space in the presence of linear degradation. In this geometry:

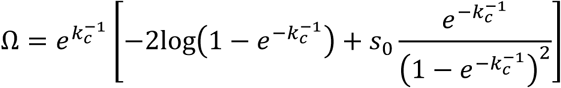

## Notes

#### Summary of Updates

Thre is a minor correction in the abstract. This was done just to synchronize the version with a journal submission

https://github.com/alesposito/CloE-PE

